# Humidity Reduces Rapid and Distant Airborne Dispersal of Viable Viral Particles in Classroom Settings

**DOI:** 10.1101/2021.06.22.449435

**Authors:** Antun Skanata, Fabrizio Spagnolo, Molly Metz, Davida S. Smyth, John J. Dennehy

**Author notes:** co-first authors (Antun Skanata, Fabrizio Spagnolo). co-corresponding authors, Address correspondence to John J. Dennehy, Biology Department, Queens College, 65-30 Kissena Boulevard, Flushing, NY 11367, USA, and Davida S. Smyth, The New School, 63 5th Avenue, Room 618A, NY 10003.

## Abstract

The transmission of airborne pathogens via aerosols is considered to be the main route through which a number of known and emerging respiratory diseases infect their hosts. It is therefore essential to quantify airborne transmission in closed spaces and determine the recommendations that should be implemented to minimize exposure to pathogens in built environments. We have developed a method to detect viable virus particles from aerosols by using an aerosolized bacteriophage Phi6 in combination with its host *Pseudomonas phaseolicola*, which when seeded on agar plates acts as a virus detector that can be placed at a range of distances away from an aerosol-generating source. Based on this method we present two striking results. (1) We consistently detected viable phage particles at distances of up to 18 feet away from the source within 15-minutes of exposure in a classroom equipped with a state of the art HVAC system. (2) Increasing the relative humidity beyond 40% significantly reduces dispersal. Our method can be used to quantify the exposure to pathogens at various distances from the source for different amounts of time, data which can be used to set safety standards for room capacity and to ascertain the efficacy of interventions that aim to reduce pathogen levels in closed spaces of specified sizes and intended uses.

**Summary:** We present a method to experimentally determine the exposure to airborne pathogens in closed spaces.

## Introduction

Airborne transmission of human pathogens has been a driver of major outbreaks of known and novel respiratory diseases ^1^. It is now known that a number of contagious diseases can spread through a process of aerosolization, where an infected individual generates virus-carrying particles that can remain suspended in the air for long periods ^2,3^. While airborne transmission has been found to rarely occur outdoors ^4^, indoor spaces may facilitate transmission via aerosols even while physically distancing ^5^. Experiments have shown that many viruses, including SARS-CoV-2, remain infectious for long periods when suspended in aerosols ^6^. As such, it is crucial to determine the distances and timescales over which pathogens can spread indoors and ask what types of shared space usage recommendations can be implemented to minimize future outbreaks.

To parameterize viral dispersal in the built environment, we developed a method to detect viable virus particles in aerosols by using a bacterium *Pseudomonas syringae* pv *phaseolicola* genetically modified to produce LacZ-***α***, which serves as the host for a LacZ-***β***-marked bacteriophage, Phi6. Phi6 is a lipid-coated icosahedral bacteriophage commonly used as a proxy for human enveloped viruses, including SARS-CoV-2, due to its similar structure, size, and physiology ^7–9^. When plated on agar containing X-Gal and the LacZ-***α*** expressing host, these phages produce easily identifiable blue plaques, which are lesions in the bacterial lawn where phages have lysed their hosts (Fig 1C, inset). X-Gal agar plates overlaid with soft agar containing the *P. phaseolicola* host and exposed for varying durations thus act as virus detectors, which can be placed at a range of distances from the aerosol-generating source of phage Phi6 (Fig 1B).

**Figure 1:**
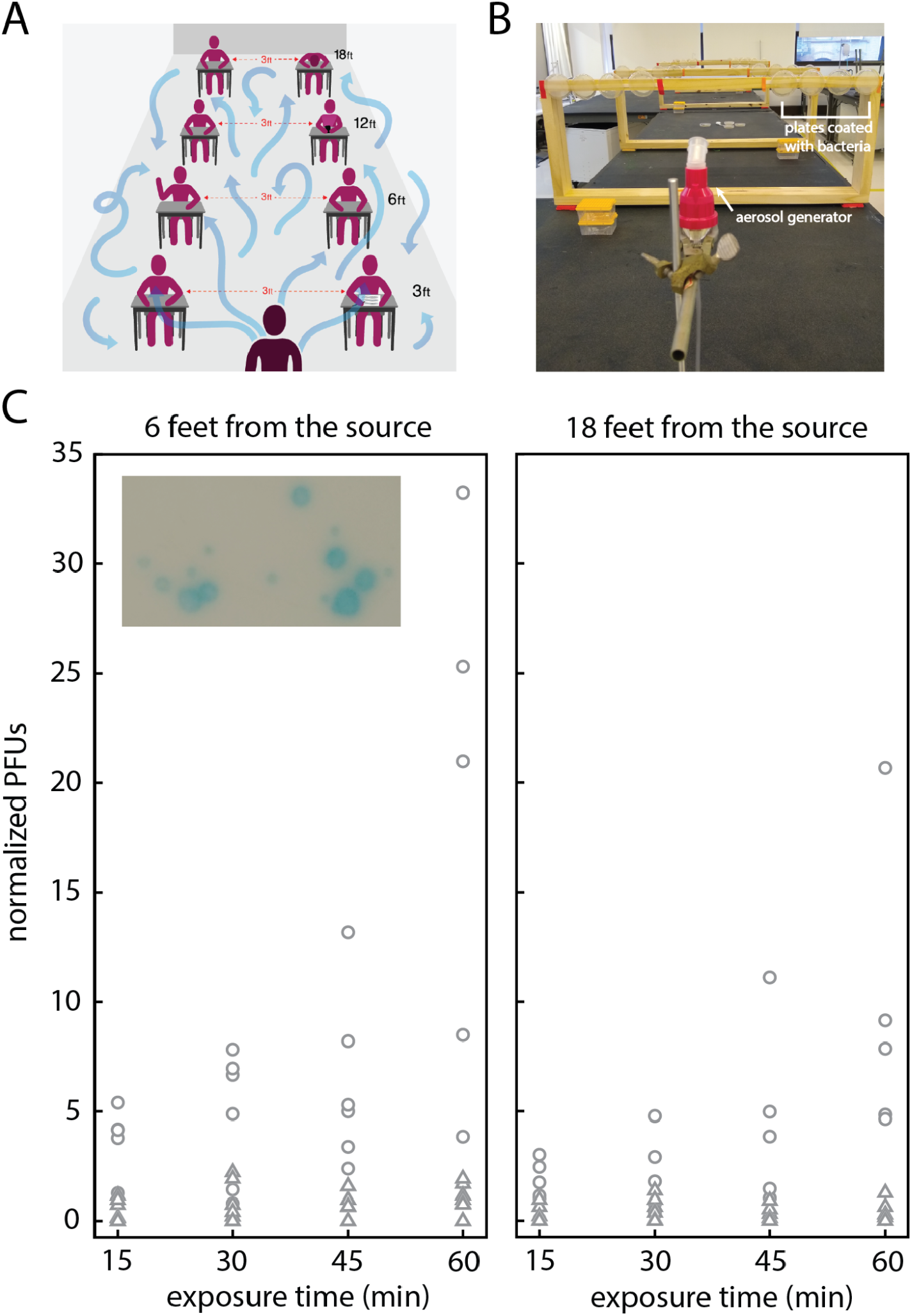
Experiments to determine the spreading of viable aerosolized pathogens in closed spaces. (A) Graphical representation of a typical classroom setting where physical distancing is enforced. Blue arrows depict the routes by which aerosolized particles might travel due to diffusion and airflows. (B) Layout of the experiment. Detectors are placed at specified distances away from a nebulizer (located in the front of the photo). At each location we placed 4 detectors, which we expose for increasing durations of time: 15, 30, 45 and 60 minutes total. (C) Plaque counts as a function of exposure time for detectors placed 6 feet away (left panel) and 18 feet away from the source (right panel). Data points correspond to data collected in 2 different rooms, in a total of 7 independent experiments. Circles correspond to data collected at RH below 40%, triangles correspond to data points collected at RH above 40%. Counts are normalized to 4×10^7^ total phage released. Data at all distances is shown in Fig. S1. Inset - blue plaque morphology shows distinct size differences.

Here, we report two major results: (1) aerosolized Phi6 particles can travel distances of 18 feet in under 15 minutes in a closed room equipped with an HVAC system operating above an industry recommended level ^10^, and (2) exposure to the aerosolized virus decreases with increases in relative humidity (RH). Specifically, we find that the exposure is significantly reduced when RH is above 40% at a room temperature of (22.8 ± 0.2) °C. These results may be useful in determining the types of interventions capable of reducing or eliminating the exposure to airborne pathogens in shared closed spaces.

## Materials and Methods

### Preparation of Viral Surrogate and Host

A single colony of LacZ-***α*** producing *P. phaseolicola* was added to 10 mL of lysogeny broth (LB) and incubated for 18 h with rotary shaking (220 rpm) at 25°C ^11,12^. 500 μL of the overnight culture was added to 50 mL of fresh LB supplemented with 200 μg/mL ampicillin and incubated for 18 h with rotary shaking (200 rpm) at 25°C to provide overnight culture for soft agar overlay plates (200 μl overnight culture in 3 ml soft agar) that served as detectors for our experiments. To produce Phi6 lysate, 5 mL of stationary-phase culture was added to 200 mL fresh LB, along with 10 μL frozen phage stock when the culture reached exponential phase^13^. Following 18 h incubation with shaking, phages were isolated by filtration through 0.22-μm filters (Durapore; Millipore, Bedford, MA). Phage particles per mL were quantified via serial dilution and plating according to standard methods (Supplementary Information)^13^.

### Generation of Aerosols

To generate aerosolized viral droplets we introduced 10 mL of diluted phage lysate into a medical grade nebulizer (Uni-HEART^™^ Lo-Flo Continuous Nebulizer, Westmed Inc), which continually generates aerosols of 2-3 μm mass median aerodynamic diameter (MMAD). Aerosols of that size are known to deposit in the nose, lungs and bronchi in adults upon inhaling ^14^. Generation of aerosols in this size range has been associated with talking and coughing ^15^. The inherent variability of aerosol sizes generated by the nebulizer might effectively capture some of the variability in aerosols that are also produced by other common actions such as breathing, sneezing, exhaling, and softly talking ^15,16^.

We diluted phage lysates to concentrations of approximately 10^8^ phage particles per mL, and estimated titers used in each experiment by spot plating 10 μL aliquots at different dilutions on replicate plates. The titers we report in SI Data Tables S1, S2 and S3 are averages over 3-5 replicates. These titers led to consistently countable numbers of plaques across all distances, exposure times and external conditions. Symptomatic individuals with coronavirus and influenza virus infections can shed 10^2^ - 10^5^ viral particles over the course of 30 minutes ^17,18^. While different viruses might require a different inoculum size to successfully infect the human host, it has been shown that even a few virions of influenza A can generate new infections ^19^.

### Classroom Experiments

We arranged the placement of the source of aerosols and the detectors in such a way to consider the risks of symptomatic or asymptomatic spreading of airborne viruses in a classroom (Fig 1A), simulating a situation where mask wearing is not enforced. The detectors -- LB plates overlaid with the host strain -- were located 3, 6, 12 and 18 feet away from the nebulizer to study the impact of physical distancing on transmission in closed spaces. We placed two sets of four detectors that are laterally 3 feet apart (Fig 1B) while being approximately the same distance away from the source. Single detectors within each set were exposed to the aerosolized phage for progressively longer durations of time for up to one hour (i.e., 15, 30, 45, & 60 minutes). Such placement of the detectors provides a replicate measurement within the same experiment. We repeated our experiments over multiple days in two different rooms at The New School, located on different floors, and whose climates are controlled by separate, independent HVAC systems. The results that we report here are consistent among the two rooms despite the variation generated by changes in external conditions, differences in room size, room layouts, and other sources (Supplementary Information & Fig. S2).

## Results

### Aerosolized phage can spread over large distances

We performed our experiments in two replicate classrooms at The New School in New York City, both of which are equipped with a state of the art HVAC system with filtration corresponding to two rows of filters (pre-filter M8 and final filter M14), which continuously operates at about 15 room air changes per hour (private correspondence). Yet, we consistently observed plaque-forming units (PFUs) on plates at distances of up to 18 feet away from the nebulizer and exposed for a duration of 15 minutes from the start of aerosol generation (Fig 1C). We found that the exposure to aerosolized phage particles at 18 feet away from the source shows on average a modest 1.6x reduction when compared to the exposure observed at 6 feet over the same durations (Fig. 1C). Therefore, we consider it likely that the airflows present inside the room facilitate the spread of aerosolized phage throughout the entire room. While it is possible that the HVAC system filters some of the phage from the environment, it most likely does so at timescales that are longer than the time it takes the aerosolized phage to spread throughout the room. As such, our experiments suggest that viable aerosolized viral particles can travel relatively large distances and rapidly generate successful infections.

### Humidity impacts the spread of aerosolized phage

We further found experimental evidence that the humidity impacts the spread and therefore the exposure to aerosolized phage. Fig. 1C shows the data we collected across two rooms and a range of external air conditions. Further inspection revealed that the air temperature was consistently maintained by the HVAC system in the range 22.5 - 23 °C across all experiments, with temperatures fluctuating on average below 1% (Fig. S3 & S4). However, we noticed an apparent difference in the data when we aligned the PFU counts with relative humidity recordings. Circles in Fig 1C that are dispersed across a range of PFUs show normalized counts collected at RH below 40%, while triangles, which are located in a narrow band at very low PFU counts, correspond to data collected at RH higher than 40%.

Figure 2 panels A-D show the rate at which PFUs are generated per 15-minute exposure time as a function of RH and distance from the nebulizer. For plates that were exposed for longer than 15 minutes, we consider the average rate of PFU formation in a 15-minute time window. At all distances, we find that increasing RH above 40% results in a substantial decrease in exposure rates. These results are consistent among the two classrooms in which we conducted our experiments, represented with circles and squares in Fig. 2. Experimental evidence suggests similar dependence on humidity for transmission of influenza ^20–22^.

**Figure 2.**
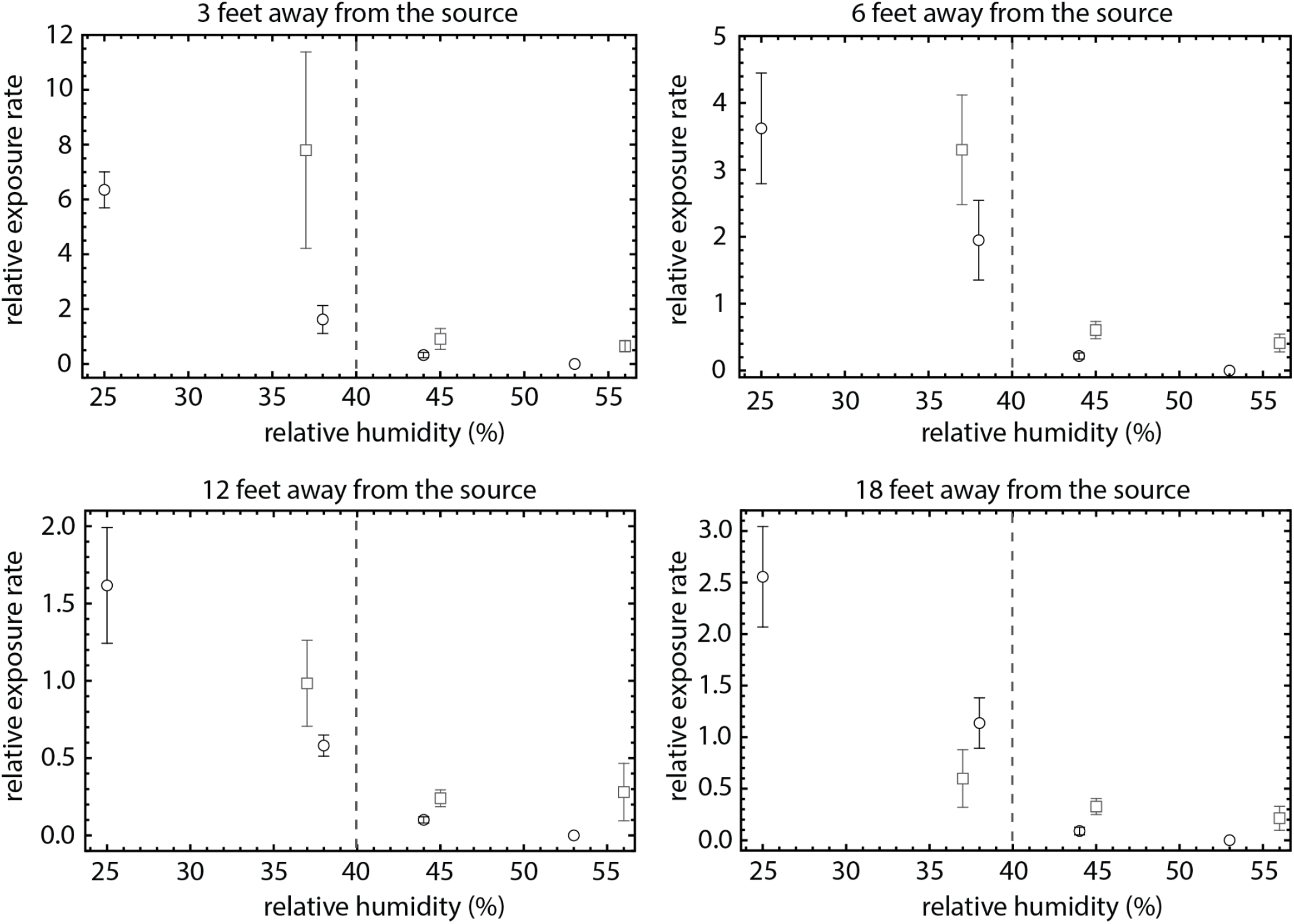
Exposure to aerosolized pathogens decreases with increasing relative humidity. Panels correspond to exposure at predetermined distances away from the source. Points are averages of PFUs normalized to 4×10^7^ total phage released and expressed as a rate per 15-minute exposure for a particular experiment, as a function of humidity, in two classrooms at The New School (rectangles -- Room 300, circles - Room 400). Error bars are ± S.E. RH has a margin of error ± 2% according to manufacturer’s specifications. Dashed vertical line at 40% relative humidity visually separates regions of high exposure (RH below 40%) and low exposure (RH above 40%).

## Discussion

Our experiments found that aerosolized phage particles can travel large distances in short amounts of time in closed spaces that are equipped with HVAC systems. We determined that the exposure to the surrogate virus significantly decreases with increases in RH. A question remains whether this decrease is a result of loss of viability at a higher humidity ^20,23^ or a result of increased surface deposition rates due to coalescing of the phage-carrying aerosols with water vapor suspended in the air ^2^. To address this question, we performed additional experiments with the detectors placed horizontally on the bench surface at distances of 3, 6,12 and 18 feet away from the nebulizer. Such placement of the detectors measures horizontal surface deposition rates at those locations. Figure S5 shows that the rate of deposition on horizontal surfaces decreases with increasing RH at distances of 3 feet and beyond, suggesting that if the surface deposition rates are increased, those increases are limited to a radius within 3 feet away from the source. The full impact of humidity and temperature on the rates of horizontal deposition is left for future work. We also performed control experiments in a room where the HVAC system was turned off (Fig. S5), and found no significant differences in exposure rates.

These results led us to speculate that, in closed spaces where levels of humidity higher than 40% cannot be achieved or controlled, a relatively inexpensive personal humidifier might provide an individualized layer of protection. We placed portable battery-operated personal humidifiers at one of the two replicate locations at different distances away from the source, and compared the exposure in the same room, under the same environmental conditions, of a set of detectors with and without a personal humidifier located at a distance of 6 in (15 cm) away from and in front of the detector plates (Fig. S6). The detectors located next to personal humidifiers showed a decrease in exposure to airborne pathogens. Experiments to further quantify the reduction in exposure are currently underway.

Observed plaques in our experiments show a range of sizes. Insertion of a genetic marker to phage Phi6 is known to induce mutations that may subsequently resolve in smaller plaque sizes ^13^. In our experiments we also noticed that plates located closer to the source (3 feet and 6 feet away) are more likely to contain larger plaques while the plates further away are more likely to contain smaller plaques, which might indicate that larger plaques may have been initially formed from larger particles that carry multiple phage. Since we cannot distinguish between a plaque generated by a single phage or multiple phage, we count each plaque as a single infection event. The true integrated exposure to aerosolized phage might be higher than our plates suggest.

We note that breathing is an active process that involves sampling the air from a volume whereas our plates detect the rates of aerosol deposition on vertical surfaces. Establishing a mapping between the counts of PFUs on plates and dosage through inhalation can be achieved by using an air sampler in parallel to the static plates and is left for future work.

Finally, we wish to comment on current recommendations that involve the use of HVAC systems coupled with physical distancing in closed spaces ^10,24^. In the case of asymptomatic transmission, we find exposure may occur at any conceivable distance, regardless of the length of time a person may be exposed. Current room capacity designations take into account space considerations, which include room surface area, placement of physical barriers, and air ventilation and filtration capabilities ^24^. Here we have shown that exposure risks to airborne pathogens in closed spaces might not be dominated by space considerations alone and that room capacity designations might be impacted by air flows and humidity. Air flows in a closed room might in turn be influenced by multiple factors, including the opening and closing of doors and windows, and the movement of people into, out of, and around the room, and are therefore dependent on the specifics of the space and the activity of persons occupying that space. Those types of considerations should be included when specific use recommendations are being made. Such efforts, which include designing ventilation systems that adjust to the number of occupants and their activity ^25^, and assessing the exposure on the basis of activity, distancing, mask wearing, and filtration efficiency ^26^, are currently underway.

The disparity between the conditions within which people of different socio-economic status live and work was a major driver of the covid-19 pandemic ^27,28^. Many office buildings and private institutions may have the funds required to implement a range of exposure-reducing interventions, while public institutions or shared-housing complexes may not. Here, we developed a method to measure the transmission of airborne pathogens in closed spaces, which is portable (can be produced in a wet lab and ported to site), relatively inexpensive, and can be deployed to a variety of spaces with different configurations and intended use. This method, supplemented with epidemiological modeling, could form a basis for inferring data-driven recommendations on room capacity and risk-reducing interventions that can be specific to the space and its intended occupants.

## Acknowledgments

We acknowledge the phage expertise, technical support and guidance of Dr. Sherin Kannoly. This work was funded by NSF Award 2032634 to Dr. John Dennehy and NSF Award 2032645 to Davida S. Smyth. We are grateful for the support of our facilities and space planning colleagues at The New School. We thank Westmed, Bob Walczer, and Ted Rozanski for help with sourcing nebulizers and compressors. We also recognize the technical support of The New School undergraduates and graduates Geena Sompanya, Simon Chen, Oriana Zwerkling and Kaitlyn Bushfield.

## Supplement

Methods text.

Fig S1. Plots of plaque counts at all distances

Fig S2. Floor plans

Fig S3 & S4. Temperature and humidity recordings

Fig S5. Surface deposition experiments

Fig S6. Experiments with personal humidifiers

Tables S1, S2 & S3: Data tables

## Supplementary Information

### Methods

#### Agar Preparation

LB medium (Miller Formula-Tryptone 10g/L, Yeast extract 5g/L, Sodium Chloride 10g/L) (BD Difco) was prepared according to the manufacturer’s instructions. LB agar plates were supplemented with 200 μg/mL 5-bromo-4-chloro-3-indolyl-β-D-galactopyranoside (X-Gal) and 200 μg/mL ampicillin (Fisher Scientific). LB was also supplemented with 200 μg/mL ampicillin (Fisher Scientific). Plates were poured the night before and kept in the dark to cool. Soft agar was prepared using LB supplemented with agar (7g per liter). 3 mL of molten soft agar was aliquoted into 13 mm glass test tubes and maintained at 48°C in a heating block. 200 μL of an overnight culture of *Pseudomonas syringae pv phaseolicola* was added, the contents vortexed gently and poured on top of an LB agar plate. For each batch of *P. phaseolicola* seeded plates, 2 plates were placed in the incubator as controls to assay for contamination.

#### Determination of Titers

Phage lysate was produced as previously described (Ref. 13 in the main text) and stored in glass bottles at 4°C until use. Before each experiment, the titer was determined using a spot method, which provides a measure of sensitivity of the detector to phage particles being deposited on its surface. Lysate was serially diluted from 10^-1^ to 10^-8^ in LB. 10 μl of each dilution was spotted onto a seeded plate. The spotted plates were incubated for two days at 25°C. The titer was determined using the formula

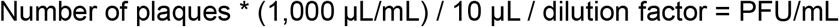

#### Choice and Calibration of Nebulizers

Several brands of commercially available nebulizers were examined for their capability to resemble what is known about simulating breathing and coughing and we chose Uni-HEART^™^ Lo-Flo Continuous Nebulizers from Westmed Inc. owing to their ability to generate aerosols in the range of 2 to 3 μm. We noticed variation in the rate of aerosolization between individual nebulizer units. We labeled all of our nebulizer units and paired them with dedicated compressors. To calibrate the rate of nebulization, we ran a number of nebulizers in parallel for one hour and periodically monitored their operation. For these runs we used 10 mL of phosphate buffered saline (PBS; Fisher Scientific) and measured the remaining volume after one hour. For our experiments we chose the nebulizers which contained similar volumes of liquid after calibration. We note the rate of nebulization depends on the media used. The nebulizers we used for the experiments expelled 5.6 ± 0.8 mL of lysate in one hour.

#### Plate placement

The plates and nebulizer were placed on benches that are 37 inches from the floor. Plates were mounted to custom-made wooden stands with double-sided mounting tape. The center of the plates was at a height of 17.5 inches (room 300) and 19 inches (room 400) above the bench owing to differences in the height of the stands used in each room. The nebulizer was held aloft using a retort stand and clamp such that the mouth opening of the nebulizer was in line with the center of the agar plates. Air ducts are located to the sides of the bench as shown in Fig. S2. Duct openings are located at a height of 6.75 ft above the surface of the bench and 29 in beneath the ceiling. The ceiling height in both rooms is 12 ft; dimensions of the respective rooms and their layouts are shown in Fig. S2. Air returns are located on the wall adjacent to the doors, approximately in line with the orientation of the benches on which experiments were done. In room 300 the nebulizer was at the opposite end of the air return and in room 400 it was below and next to the air return (see Fig. S2), which allowed us to test two orientations of the experiment with respect to the overall direction of the airflow.

#### Experimental setup

Each plate was labelled with the date, room number, distance, exposure time and location of the plate with respect to the nebulizer. While the plates were being set up, each nebulizer was started without liquid to allow residual alcohol from cleaning (see below) to evaporate. The humidity and temperature was monitored using Govee Model:H5074 devices (accuracy ± 0.2°C and ± 2% RH) placed on the bench surface at a distance 9 feet away from the nebulizer beginning 30 min before the experiments began to establish a baseline. Figures S3 and S4 show temperature and humidity charts for each individual experiment.

To start an experiment, we added 10 mL of phage lysate to the nebulizer, removed the lids from the plates and started the compressor. At the 15, 30, 45 and 60 minute marks, an experimenter entered the room, covered and removed a set of plates from the room. After all the plates were removed, the compressor was turned off.

The plates were incubated for 2-4 days at 25°C. Plaques were counted after two days of incubation after which they were moved to the bench. We did not observe additional plaques forming by the 4th day. After the experiment, the nebulizer was detached from the compressor and the remaining liquid removed using a thin tube attached to a syringe. The liquid was removed to a 15 mL conical tube and the volume was recorded. The extracted liquid was kept at room temperature for several days to monitor for contamination while the plates from the experiment were incubating. After each use, the nebulizers were washed in deionized water, the water removed, and the unit sprayed with 70% ethanol. The remaining alcohol was removed with a syringe and allowed to dry out before the next experiment. The nebulizers were replaced after 5-10 uses, after they started to leak or if they showed signs of contamination.

**Figure S1:**
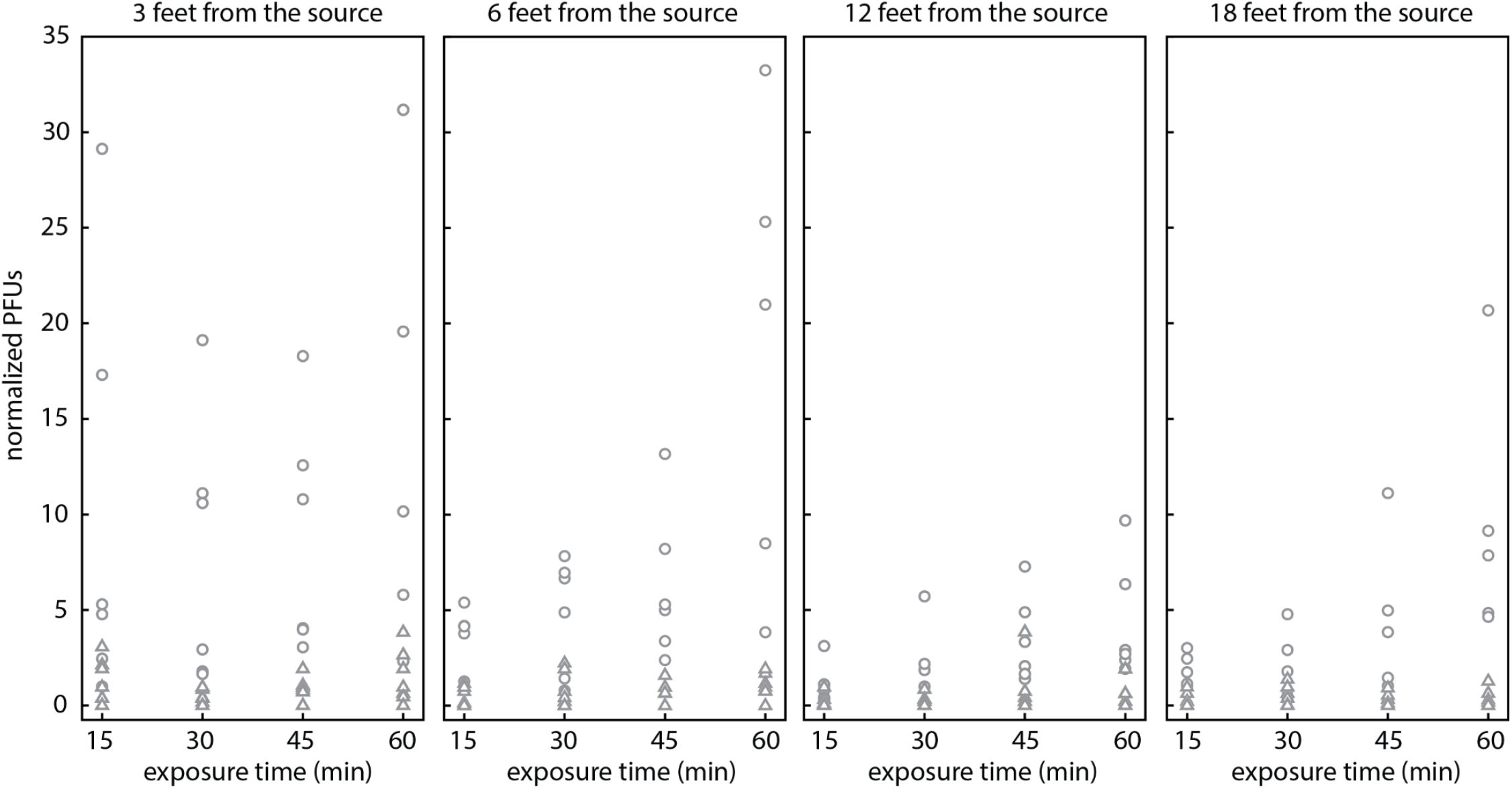
Plaque counts as a function of exposure time for detectors placed at all distances away from the source. Circles - data collected at relative humidities below 40%, triangles - data collected at relative humidities above 40%. Counts are normalized to 4×10^7^ total phage released.

**Figure S2:**
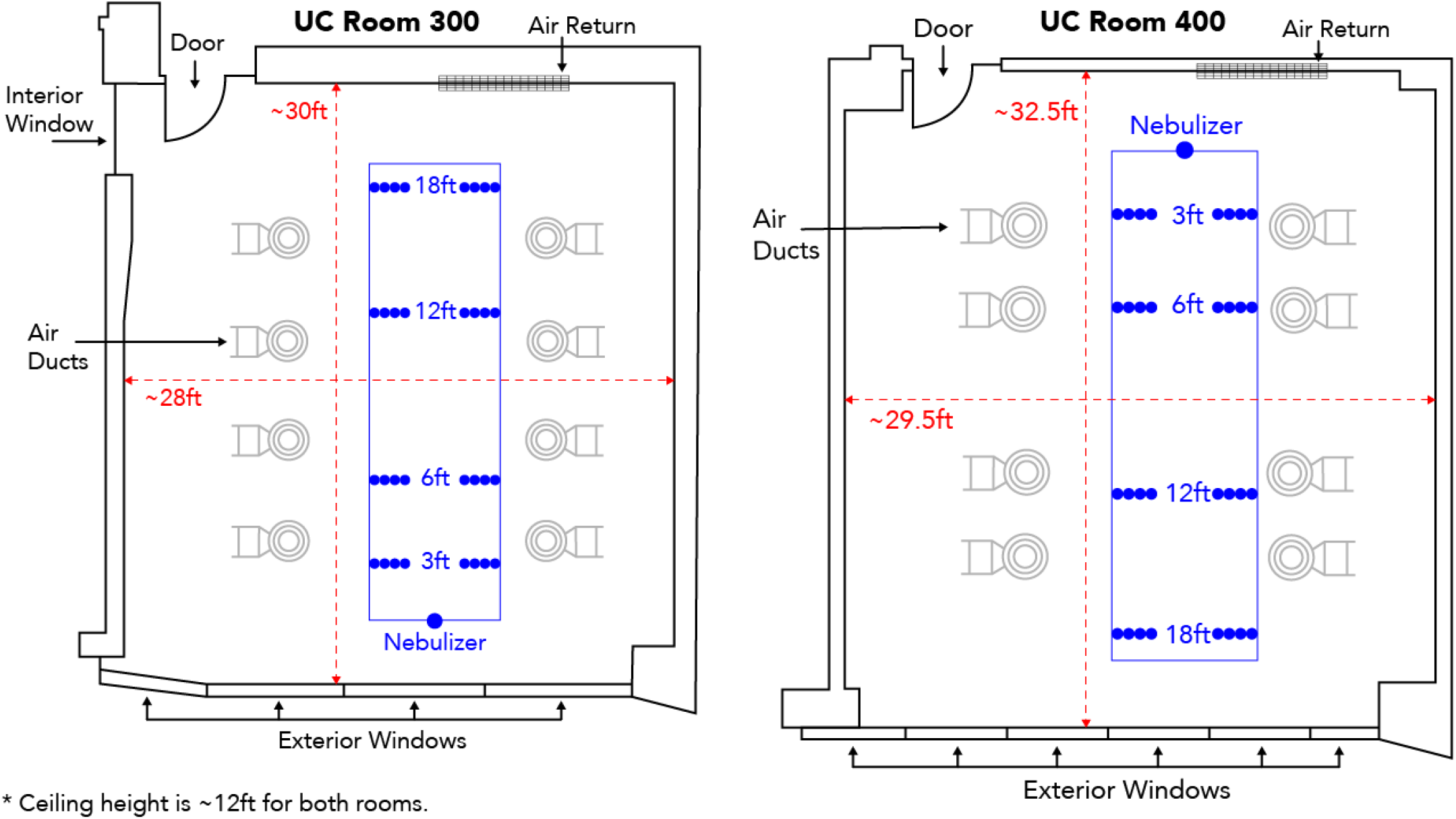
Floor plans of rooms 300 & 400 with approximate dimensions, placement of air ducts, air return, and their relation to the experimental setup.

**Figure S3:**
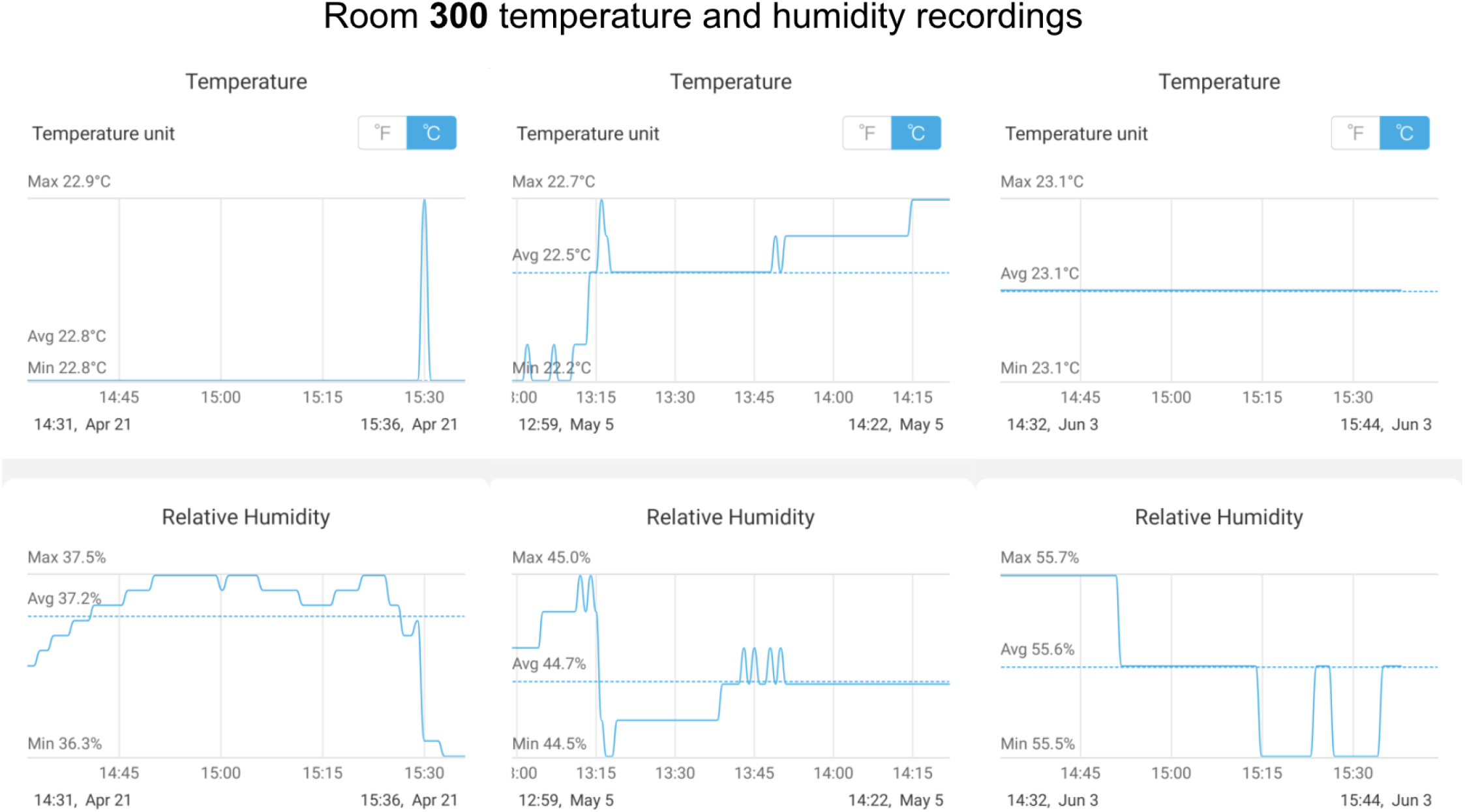
Temperature and humidity data for Room 300

**Figure S4:**
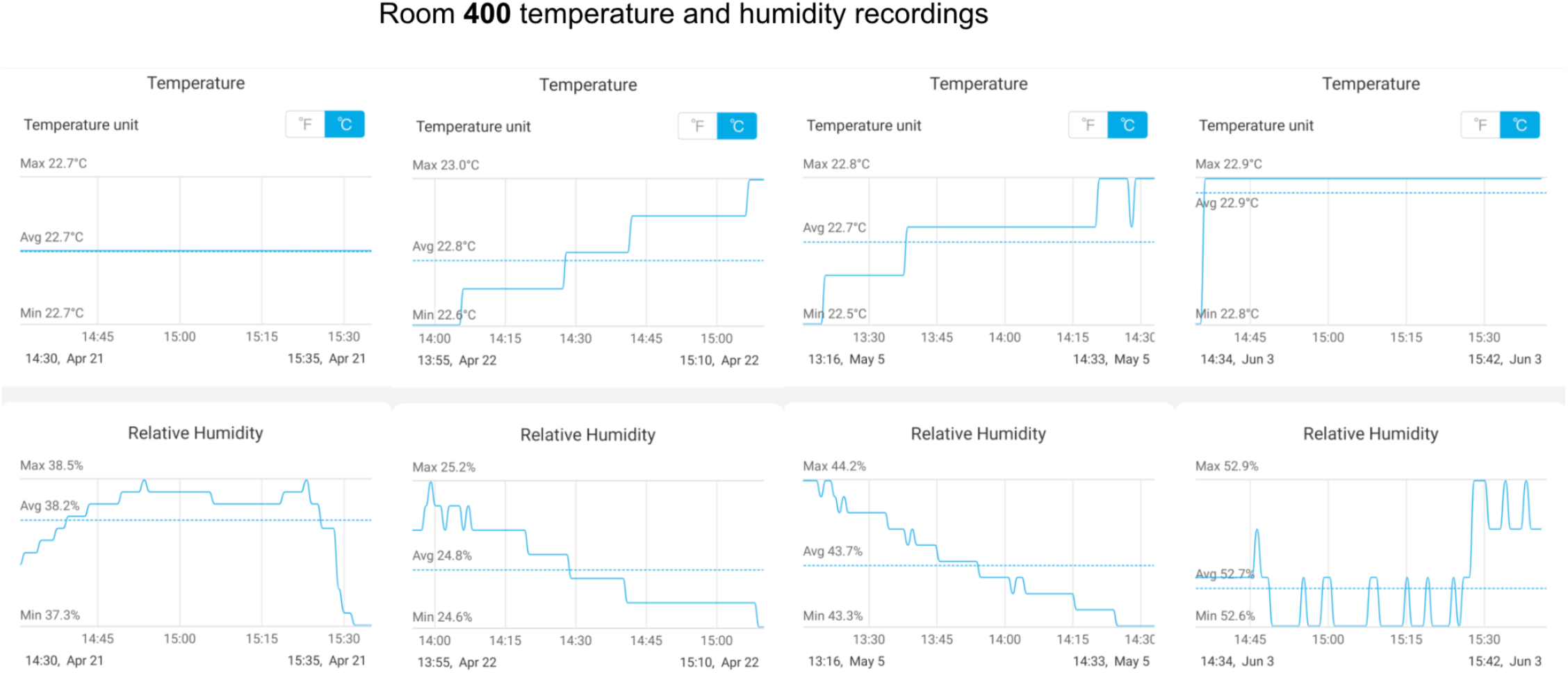
Temperature and humidity data for Room 400

**Figure S5:**
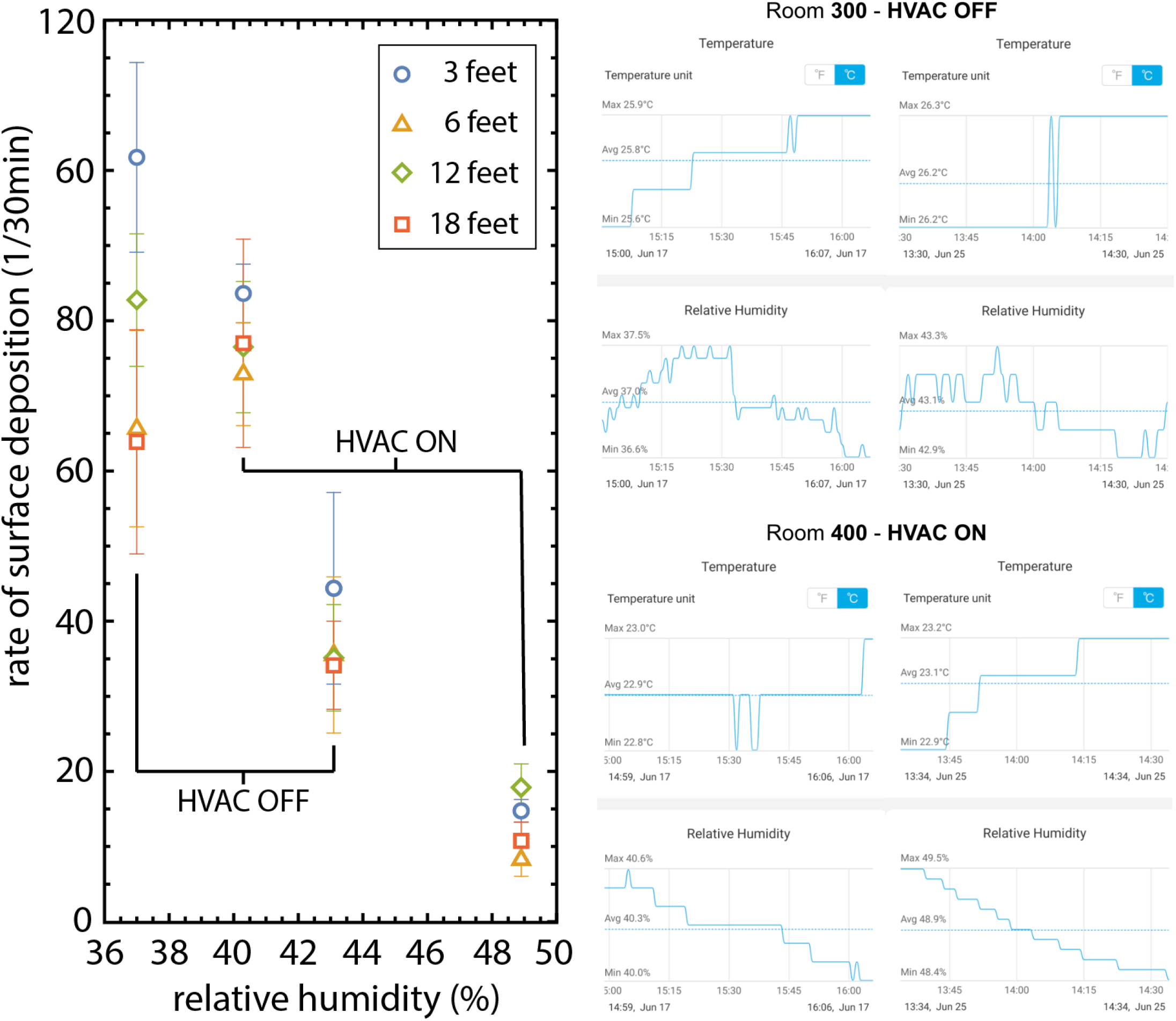
Horizontal surface deposition experiments with and without HVAC. We placed two sets of two plates horizontally on the bench surface, three feet apart, on the left and right side of the nebulizer, and a set of two vertical plates in the center. One plate in each set was exposed for 30 minutes, and the other for 60 minutes (See **Table S3**). In the room in which the HVAC was off we recorded temperatures of 25.8°C and 26.2°C, while in the room with HVAC on we recorded 22.9°C and 23.1 °C. Relative humidity ranged between 36-50% (see detailed recordings in the right panel). Data points correspond to rates of PFUs in a 30-minute time window, averaged over the two replicates and two timepoints, for 3×10^8^ total phage released. Error bars are S.E. We observe a decrease of PFUs with the increase in humidity, concurring with the results presented in the main text on vertical plates.

**Figure S6:**
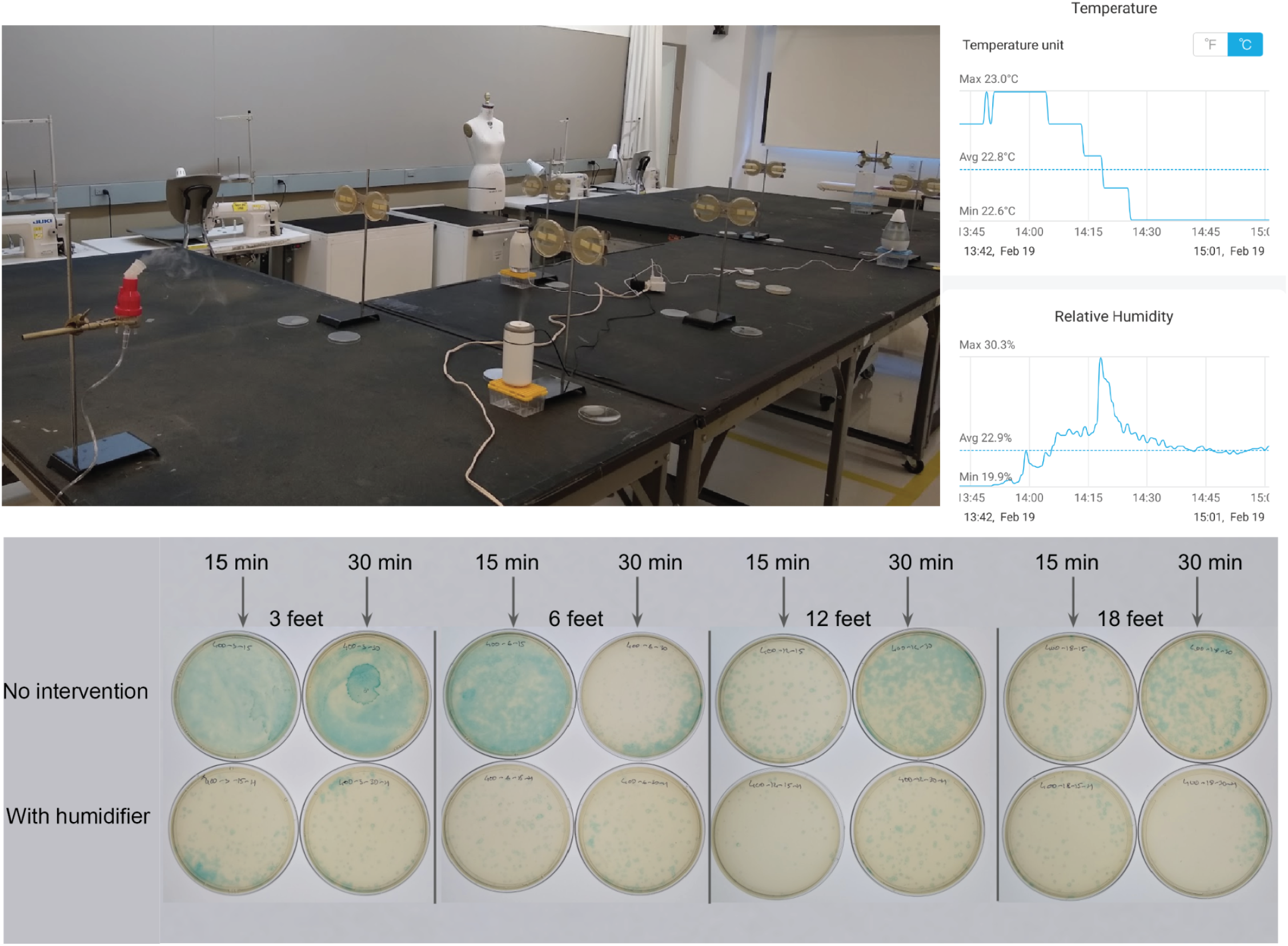
Experiments with personal humidifiers. We tested a number of personal humidifiers which we placed in front of one set of plates, while the other set at the same distance from the source had no intervention. We arranged the humidifiers in a zig-zag pattern to control for port-starboard preference. We started the humidifiers 15 minutes prior to starting phage aerosolization. We noticed a transient increase in relative humidity at a location of the measuring device, which was 9 feet away from the nebulizer (see temperature and humidity recordings in the top right panel). Plates with the humidifier show reduction of plaque counts when compared to the plates without intervention within the same experiment.

**Table S1:** PFU counts collected in **Room 300** (superscript ^C^ next to an entry means that plate had a contaminant, the number indicates how many contaminants were observed)

|  |  |  |  |  |  |  |  |  |
| --- | --- | --- | --- | --- | --- | --- | --- | --- |
| Date: April 21st, 2021 | | Titer $9 \times 10^7$ PFU/mL | | Volume nebulized: 5.6 mL | Relative Humidity: 37.2% | Average Temp.: 22.8°C | Total phage $5 \times 10^8$ | Controls: 0, 0 |
| Replicates | L | R | L | R | L | R | L | R |
| Distance/Time | 3 feet |  | 6 feet |  | 12 feet |  | 18 feet |  |
| 15 min | 218 | 367 | 16 | 68 | 14 | 11 | 22 | 1 |
| 30 min | 37 | 140 | 18 | 84 | 1 | 1 | 10 | 4 |
| 45 min | 51 | 136 | 166 | 63 | 26 | 42 | 3 | 13 |
| 60 min | 73 | 128 | 107<sup>c1</sup> | 319 | 80 | 122 | 99 | 1 |
| Date: May 5th 3rd, 2021 | | Titer $9 \times 10^7$ PFU/mL | | Volume nebulized: 4.2 mL | Relative Humidity: 44.7% | Average Temp.: 22.5°C | Total phage $4 \times 10^8$ | Controls: 0, 0 |
| Replicates | L | R | L | R | L | R | L | R |
| Distance/Time | 3 feet |  | 6 feet |  | 12 feet |  | 18 feet |  |
| 15 min | 20 | 29 | 11 | 7 | 3 | 1 | 6 | 2 |
| 30 min | 4 | 8 | 7 | 21 | 3 | 8 | 6 | 13 |
| 45 min | 10 | 9 | 6 | 15 | 4 | 7 | 3 | 5 |
| 60 min | 6 | 25 | 16 | 11 | 2 | 18 | 6 | 12 |
| Date: June 3rd, 2021 | | Titer $8 \times 10^6$ PFU/mL | | Volume nebulized: 5.2 mL | Relative Humidity: 55.6% | Average Temp.: 23.1°C | Total phage $4 \times 10^7$ | Controls: 0, 0 |
| Replicates | L | R | L | R | L | R | L | R |
| Distance/Time | 3 feet |  | 6 feet |  | 12 feet |  | 18 feet |  |
| 15 min | 2 | 1 | 1 | 0 | 0 | 1 | 1 | 0 |
| 30 min | 1 | 1 | 2 | 0 | 0 | 0 | 1 | 1 |
| 45 min | 0 | 2 | 1 | 1 | 4 | 0 | 0 | 0 |
| 60 min | 1 | 2 | 1 | 2 | 0 | 0 | 0 | 0 |

**Table S2:**
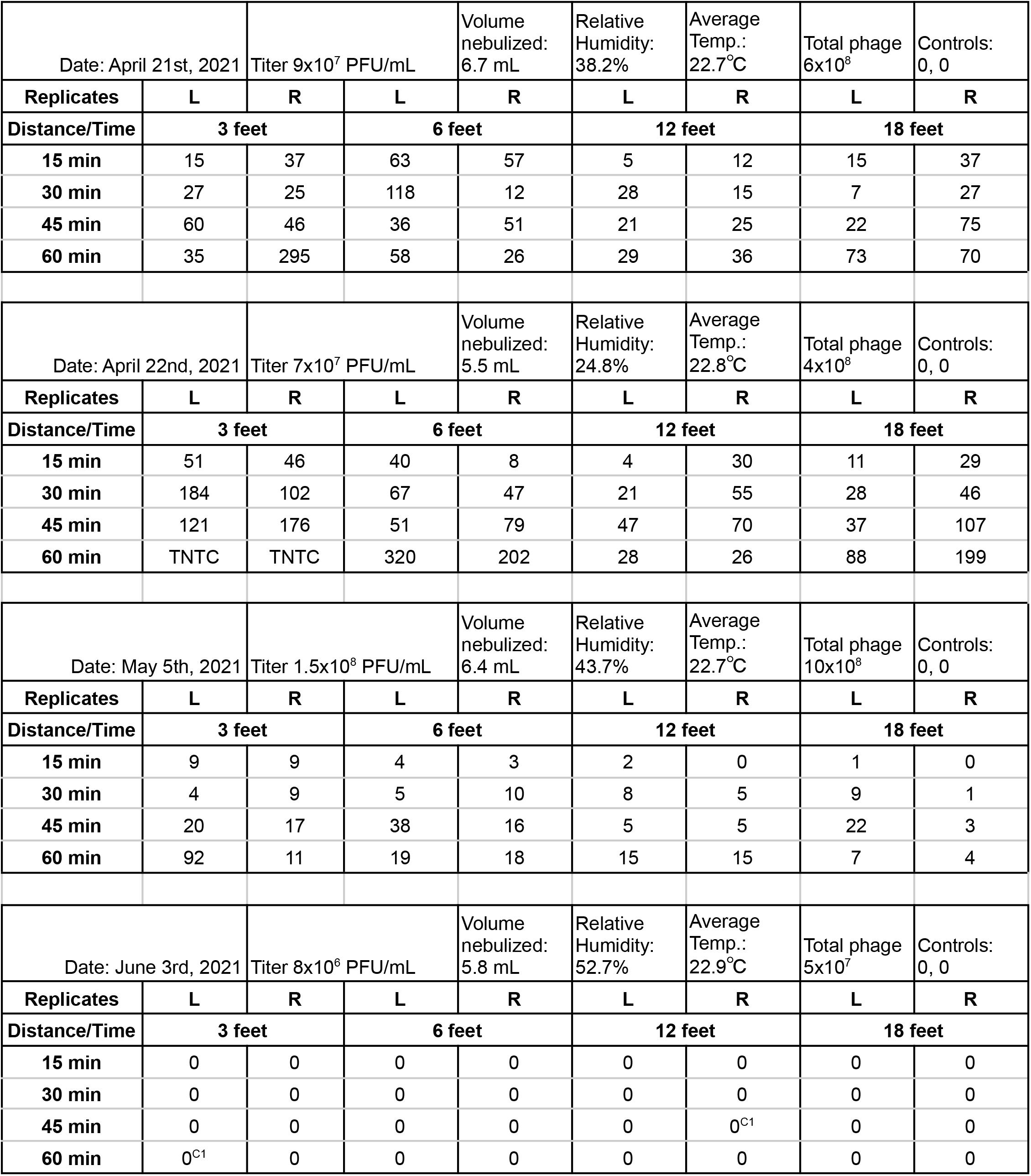
PFU counts collected in **Room 400** (superscript ^C^ next to an entry means that plate had a contaminant; TNTC - too numerous to count -- in those cases we used 300 as a reference value)

**Table S3:** PFU counts collected in surface deposition experiments in room 300 while the HVAC was off and in room 400 while the HVAC was on. (L - horizontal plates on the port side, R - horizontal plates on the starboard side, V - vertical plates in the center)

|  |  |  |  |  |  |  |  |  |  |  |  |  |
| --- | --- | --- | --- | --- | --- | --- | --- | --- | --- | --- | --- | --- |
| **Room 400**<br>**HVAC ON** | Date: June 17th, 2021 | | Titer $5 \times 10^7$ PFU/mL | | Volume nebulized: 6mL | | RH: 40.3%, T: 22.9°C | | | Total phage $3 \times 10^8$ | | Controls: 0, 0 |
| **Plate placement** | **L** | **V** | **R** | **L** | **V** | **R** | **L** | **V** | **R** | **L** | **V** | **R** |
| **Distance/Time** | **3** |  |  | **6** |  |  | **12** |  |  | **18** |  |  |
| **30** | 76 | 1 | 92 | 73 | 1 | 72 | 95 | 2 | 73 | 64 | 24 | 49 |
| **60** | 177<sup>C1</sup> | 9 | 156 | 180 | 9 | 113 | 168 | 27 | 108 | 227 | 63 | 163 |
| **Room 300**<br>**HVAC OFF** | Date: June 17th, 2021 | | Titer $5 \times 10^7$ PFU/mL | | Volume nebulized: 5.6mL | | RH: 37%, T: 25.8°C | | | Total phage $3 \times 10^8$ | | Controls: 0, 0 |
| **Plate placement** | **L** | **V** | **R** | **L** | **V** | **R** | **L** | **V** | **R** | **L** | **V** | **R** |
| **Distance/Time** | **3** |  |  | **6** |  |  | **12** |  |  | **18** |  |  |
| **30** | 115 | 34 | 65<sup>C1</sup> | 34 | 8 | 55 | 61 | 9 | 103<sup>C1</sup> | 66 | 20 | 26 |
| **60** | 242 | 12 | 212<sup>C1</sup> | 164 | 34 | 183 | 177 | 15 | 157 | 129<sup>C1</sup> | 85 | 198<sup>C1</sup> |
| **Room 400**<br>**HVAC ON** | Date: June 25th, 2021 | | Titer $4 \times 10^7$ PFU/mL | | Volume nebulized: 7mL | | RH: 48.9%, T: 23.1°C | | | Total phage $3 \times 10^8$ | | Controls: 0, 0 |
| **Plate placement** | **L** | **V** | **R** | **L** | **V** | **R** | **L** | **V** | **R** | **L** | **V** | **R** |
| **Distance/Time** | **3** |  |  | **6** |  |  | **12** |  |  | **18** |  |  |
| **30** | 16 | 7 | 11 | 13 | 1 | 4 | 16 | 1 | 13 | 17 | 1 | 6 |
| **60** | 36 | 3 | 28 | 10 | 2 | 22 | 31 | 0 | 54<sup>C1</sup> | 15 | 5 | 25 |
| **Room 300**<br>**HVAC OFF** | Date: June 25th, 2021 | | Titer $4 \times 10^7$ PFU/mL | | Volume nebulized: 6.8mL | | RH: 43.1%, T: 26.2°C | | | Total phage $3 \times 10^8$ | | Controls: 0, 0 |
| **Plate placement** | **L** | **V** | **R** | **L** | **V** | **R** | **L** | **V** | **R** | **L** | **V** | **R** |
| **Distance/Time** | **3** |  |  | **6** |  |  | **12** |  |  | **18** |  |  |
| **30** | 14 | 30 | 36 | 28 | 1 | 11 | 18 | 3 | 33 | 43 | 4 | 20 |
| **60** | 148 | 20 | 107 | 87 | 3 | 119 | 74 | 63 | 105 | 58 | 38 | 89 |

## Notes

### Competing Interest Statement

The authors have declared no competing interest.

